# *Staphylococcus aureus* biofilm secreted factors cause mucosal damage, mast cell infiltration and goblet cell hyperplasia in a rat rhinosinusitis model

**DOI:** 10.1101/2023.03.29.534842

**Authors:** Ghais Houtak, Roshan Nepal, George Bouras, Gohar Shaghayegh, Catherine Bennett, John Finnie, Kevin Fenix, Alkis James Psaltis, Peter-John Wormald, Sarah Vreugde

## Abstract

Chronic Rhinosinusitis (CRS) is an inflammatory condition of the paranasal sinus mucosa. Despite being a common health issue, the exact cause of CRS is yet to be understood. However, research suggests that *Staphylococcus aureus*, particularly in the biofilm form, drives the disease. This study aimed to investigate the impact of long-term exposure to secreted factors of *Staphylococcus aureus* biofilm (SABSF), harvested from clinical isolates of non-CRS carriers and CRS patients, on the nasal mucosa in a rat model.

Wistar rats were randomised (n=5/group) to receive daily intranasal instillations of 40 μL (200 μg/μL) SABSF for 28 days or vehicle control with *S. aureus* isolated from the sinuses of a non-CRS carrier, a type 2 endotype CRS with nasal polyps (CRSwNP) patient, and a non-type 2 endotype CRS without nasal polyps (CRSsNP) patient. The sinonasal samples of the rats were then analysed through histopathology and transcriptome profiling.

The results showed that all three intervention groups displayed significant lymphocytic infiltration (p≤0.05). However, only the SABSF collected from the CRSwNP patient caused significant mucosal damage, mast cell infiltration, and goblet cell hyperplasia compared to the control. The transcriptomics results indicated that SABSF significantly enriched multiple inflammatory pathways and showed distinct transcriptional expression differences between the control group and the SABSF collected from CRS patients (p≤0.05). Additionally, the SABSF challenges induced the expression of IgA and IgG but not IgE.

In conclusion, this *in vivo* study indicates that long-term exposure to SABSF leads to an inflammatory response in the nasal mucosa with increased severity for *S. aureus* isolated from a CRSwNP patient. The findings of this study shed light on the role of *S. aureus* in the development of CRS and could inform future research and treatment efforts.

## Introduction

Chronic rhinosinusitis (CRS) is a chronic inflammatory disease involving the mucosal lining of the nasal passage and paranasal sinuses, affecting 10-15% of the Western population (Fokkens et al., 2020; Hastan et al., 2011). The inflammatory microenvironment in the sinonasal tissue of CRS patients comprises a heterogeneous mixture of inflammatory cells that can be polarised towards type 1, 2 or 3 immunity (Wang et al., 2016). CRS patients with nasal polyps (CRSwNP) predominantly have a type 2 (T2) immune polarisation with tissue eosinophilia and increased IL-5, IL-4 and Il-13 cytokine levels. In contrast, most CRS without nasal polyps (CRSsNP) patients do not show inflammatory polarisation and are therefore classified as non-T2 endotypes, which are often accompanied by increased neutrophil infiltration in the mucosa (Grayson, Cavada, & Harvey, 2019; Kato, Peters, et al., 2022). The pathophysiology of CRS cannot be attributed to a single factor, but rather, multiple host intrinsic and environmental factors are thought to play a role (Fokkens et al., 2020).

Numerous studies have associated *Staphylococcus aureus* as a potential driver of disease in CRS (Vickery, Ramakrishnan, & Suh, 2019). Indeed, *S. aureus* is often cultured from the sinuses of CRS patients during the exacerbation of the disease (Okifo, Ray, & Gudis, 2022). Furthermore, *S. aureus* induces secretion of various cytokines such as thymic stromal lymphopoietin (TSLP), IL-33 and IL-22 in nasal mucosa (Lan et al., 2018; Mulcahy, Leech, Renauld, Mills, & McLoughlin, 2016). These cytokines activate immune orchestrating cells such as group 2 innate lymphoid cells and T-helper (Th) 2 cells, which in turn drive type 2 immunity (Miljkovic et al., 2014; Stanbery, Shuchi, Jakob von, Tait Wojno, & Ziegler, 2022). Another mechanism in which *S. aureus* is thought to be involved in the pathophysiology of CRS is mediated by IgE antibodies targeting *S. aureus* enterotoxins (Bachert, Gevaert, Holtappels, Johansson, & van Cauwenberge, 2001; Van Zele et al., 2004). It has been shown that increased IgE contributes to the acute and chronic symptoms of allergic airway disease (Teufelberger, Broker, Krysko, Bachert, & Krysko, 2019). Among CRS patients, *S. aureus* enterotoxin–specific IgEs are predominantly seen in CRSwNP patients (Tomassen et al., 2016). Multiple lines of research have suggested that the *S. aureus* enterotoxin–specific IgEs found in CRS tissue is locally secreted by plasma cells (Gevaert et al., 2013; Takeda et al., 2019; Van Zele et al., 2006).

*S. aureus* is commonly found as a commensal bacterium in the human nares and is highly adaptable to its environment (Malachowa & DeLeo, 2010; Tong, Davis, Eichenberger, Holland, & Fowler, 2015). Although the co-culturing of *S. aureus* and respiratory epithelial cells results in the elevation of inflammatory markers *in vitro*, few studies have investigated the exposure of nasal epithelium to *S. aureus* biofilm-secreted factors *in vivo*. Furthermore, the impact of disease-specific *S. aureus* strains on the nasal mucosa is unclear. Lastly, much uncertainty still exists about the mechanisms that underpin the local secretion of *S. aureus* enterotoxin–specific IgE in the nasal mucosa.

Therefore, this study aimed to elucidate the effect of long-term exposure to secreted factors of *S. aureus* strains harvested from CRS patients and controls grown in biofilm form.

## Methods

### Ethics

This study complies with all relevant ethical regulations for animal and human research. Ethical approval for animal experimentation was obtained from The University of Adelaide’s Animal Ethics Committee (M-2019-101) and conducted according to the Australian code for the care and use of animals for scientific purposes (Hubrecht, 2013). Ethics approval for isolation, storage and use of clinical isolates and tissue samples of CRS subjects was granted by The Central Adelaide Local Health Network Human Research Ethics Committee (reference HREC/15/TQEH/132).

### *S. aureus* clinical isolates

Three *S. aureus* clinical isolates (CIs) were selected from our bacterial biobank comprised of samples collected from ear-nose-throat inpatient and outpatient clinics. The CIs were selected based on the host subject’s disease phenotype and inflammatory pattern in the sinus tissue. Two CIs were from CRS patients; one CRSwNP with a type 2 inflammatory endotype and one CRSsNP with a non-type 2 endotype (Grayson et al., 2019). One CI was isolated from a non-CRS control subject with no evidence of sinonasal inflammation based on the absence of CRS symptoms and endoscopic evaluation. The inflammatory status of the tissues was assessed using flow cytometry. In brief, fresh sinonasal tissue samples underwent enzymatic digestion with a mixture of 25mg/mL collagenase D (Roche Diagnostics GmbH, 11088858001, Mannheim, Germany), 10mg/mL DNAse I (Sigma-Aldrich, 11284932001, St. Louis, USA), and Hanks’ Balanced Salt Solution (Thermo Fisher Scientific, 88284, Waltham, USA) for 30 minutes at 37°C. The resulting single-cell suspensions were used at a concentration of 4 million cells/mL and stained with Fixable Viability Dye eFluor 780 (BD Bioscience, 565388, San Jose, USA), followed by Fc Block incubation and labelling with fluorochrome-conjugated antibodies (CD45-PerCP-Cy5.5, CD14-Alexa-Flour 488, CD16-BV510, and CD24-PE). Multi-colour flow cytometry was performed using a BD FACS Canto II instrument (BD Bioscience) and FACS Diva software, with at least 500,000 events collected per sample. Data analysis was performed using FlowJo software v10.8.1.

### *S. aureus* biofilm-secreted factors

SABSF were used as stimulants. The SABSF were collected from the supernatant of *S. aureus* after biofilm formation. In brief, the CIs were cultured on Nutrient Agar (Sigma-Aldrich, N7519, USA). Then, single colonies were suspended in 0.9 % saline to a 1 McFarland Unit (MFU) turbidity reading. The suspension was diluted 15-fold in Nutrient Broth before incubation in a 6-well microtiter plate (2 mL per well). The inoculated suspension was cultured to induce biofilm formations; the microtiter plates were incubated for 48 hours at 37°C with sheer force on a rotating plate set at 70 rpm (Ratek Instruments, 3D Gyratory Mixer, Boronia, Australia). Following the incubation, the biofilm was dispersed by pipetting (minimum of 10 times), and the bacterial broth culture was collected and centrifuged for 10 mins at 1500 relative centrifugal force (rcf) in 4°C. The supernatants were sterilised through a 0.22 µm acrodisc filter (Pall Corporation, 4612, Port Washinton, USA). The filtered supernatants were concentrated 8-fold using a pre-rinsed 3K MWCO Pierce Protein Concentrator PES (molecular-weight cutoff: 3 kDa) (Thermo Fisher Scientific, 88525, USA) by centrifuging at 4000 rcf in 4°C, according to the manufacturer’s instructions. The protein concentration of the concentrated supernatants was quantified using NanoOrange Protein Quantitation Kit (Thermo Fisher Scientific, N6666). The concentrated supernatants, or SABSF, were diluted with Nutrient Broth to a concentration of 200 µg/µL and stored at -80°C until further use.

### Animals and study design

Twenty male Wistar rats (Animal Resources Centre, Perth, Australia) were obtained at 11 to 12 weeks of age. Animals were then randomised into four treatment groups receiving either (1) intranasal challenges of SABSF harvested from a *S. aureus* strain isolated from a T2 endotype CRSwNP patient, (2) a non-T2 endotype CRSsNP patient, (3) a non-CRS *S. aureus* nasal carrier, or (4) vehicle-control solution (n = 5 per group). Using a short induction with isoflurane anaesthetic gas, the SABSF (200 µg/µL) was administered as 40 μL intranasally (20 μL in each nare) daily for 28 days. On day 28, the animals were sacrificed, and the nasal tissue was harvested for histological and transcriptomics analysis (Figure 1).

**Figure 1.**
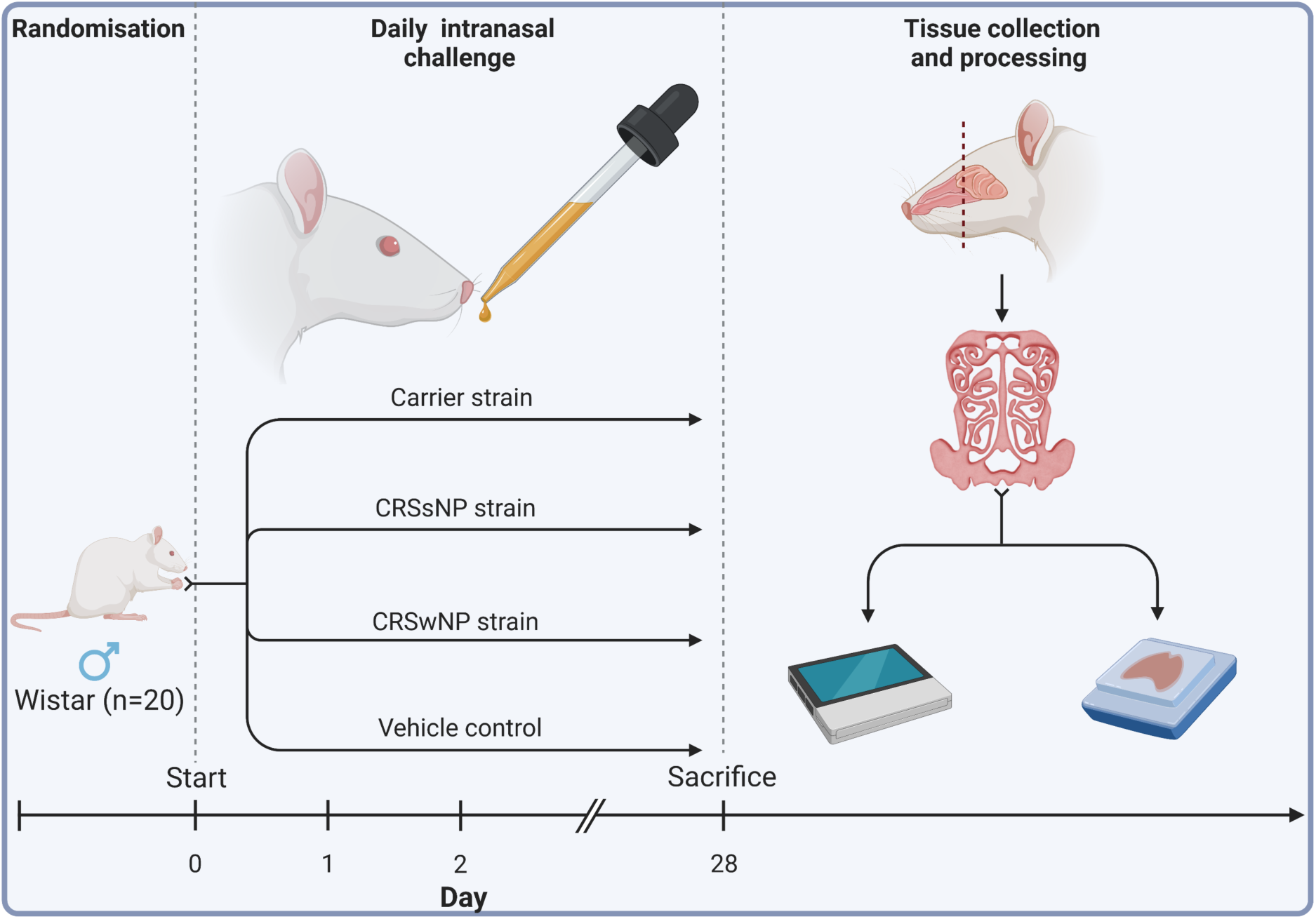
Schematic outline of the experimental design. Twenty male Wistar rats were randomised into four groups (n = 5 per group). The animals received intranasal challenges of secreted factors of the *S. aureus* biofilms with strains isolated from a CRSwNP patient, a CRSsNP patient, a non-CRS carrier, or the vehicle-control solution. Each animal received 40 μL of SABSF (200 µg/µL) intranasally (20 μL in each nare) daily for 28 days, after which the nasal tissue was harvested for histology and transcriptomics. Tissue samples of all animals were processed for histological analysis. Three animals per group were processed for long-read direct cDNA transcriptomics on the Oxford Nanopore Technologies (ONT) platform. Created with biorender.com.

### Tissue collection

The heads of the animals were collected, and complete transverse sections of the sinonasal structures were cut at 0.5 mm rostral to the anterior margin of the orbit. The rostral section was placed overnight in RNAlater Stabilisation Solution (Thermo Fisher Scientific, AM7020) and stored at −80 °C until RNA extraction, while the caudal sections were fixed in 10% neutral-buffered formalin for 48 hours at RT.

### Histological staining

After formalin fixation, the samples were decalcified in 10% ethylenediaminetetracetic acid (EDTA, pH:7.0) (Sigma-Aldrich, RDD017) solution for two weeks, paraffin-embedded, and 4µm sections cut and stained with haematoxylin and eosin (H&E). Duplicate sections were stained with periodic acid-Schiff (PAS) and toluidine blue to detect goblet and mast cells. In order to detect T-cell infiltration of mucous membranes, sections were immunostained for CD3, a specific marker for T-cell derivation. For CD3 immunohistochemistry, samples underwent heat-induced antigen retrieval in the Aptum Bio Retriever 2100 (Diagnostic technology, Belrose, Australia) using 10 mM sodium citrate buffer (pH: 6). The sections were incubated overnight with anti-cell marker cluster of differentiation (CD) 3 antibody (1:200, Abcam, ab16669, Cambridge, UK) at 4°C in a humidified incubation chamber. Afterwards, the sections were incubated in Cy-5 anti-rabbit secondary antibody (1:1000, Jackson ImmunoResearch Labs Inc., AB_2340607, West Grove, USA) for 1 hour at 4°C and counterstained with 4’,6-Diamidino-2-Phenylindole, Dihydrochloride (DAPI) (Thermo Fisher Scientific, D1306). A negative control omitting the primary antibody and a positive control showing the typical pattern of expression of this antigen were run with each batch of slides. No-primary-antibody controls were used to determine background staining intensities.

### Histological analysis

All sections were scanned using the Hamamatsu Photonics Digital Slide Scanner at 40x magnification (NanoZoomer S60, Hamamatsu, Japan). The IHC-IF stained slides were scanned on the Zeiss Axio Scan Z1 Slide Scanner (Carl Zeiss Microscopy, Oberkochen, Germany) at a magnification of 40x. Images were imported into the open-source digital pathology software QuPath for analysis (Bankhead et al., 2017).

Goblet cell hyperplasia, epithelial ulceration, and mast cell infiltration were scored using a semi-quantitative scale (Table 1)(Herbert, Janardhan, Pandiri, Cesta, & Miller, 2018).

**Table 1.**
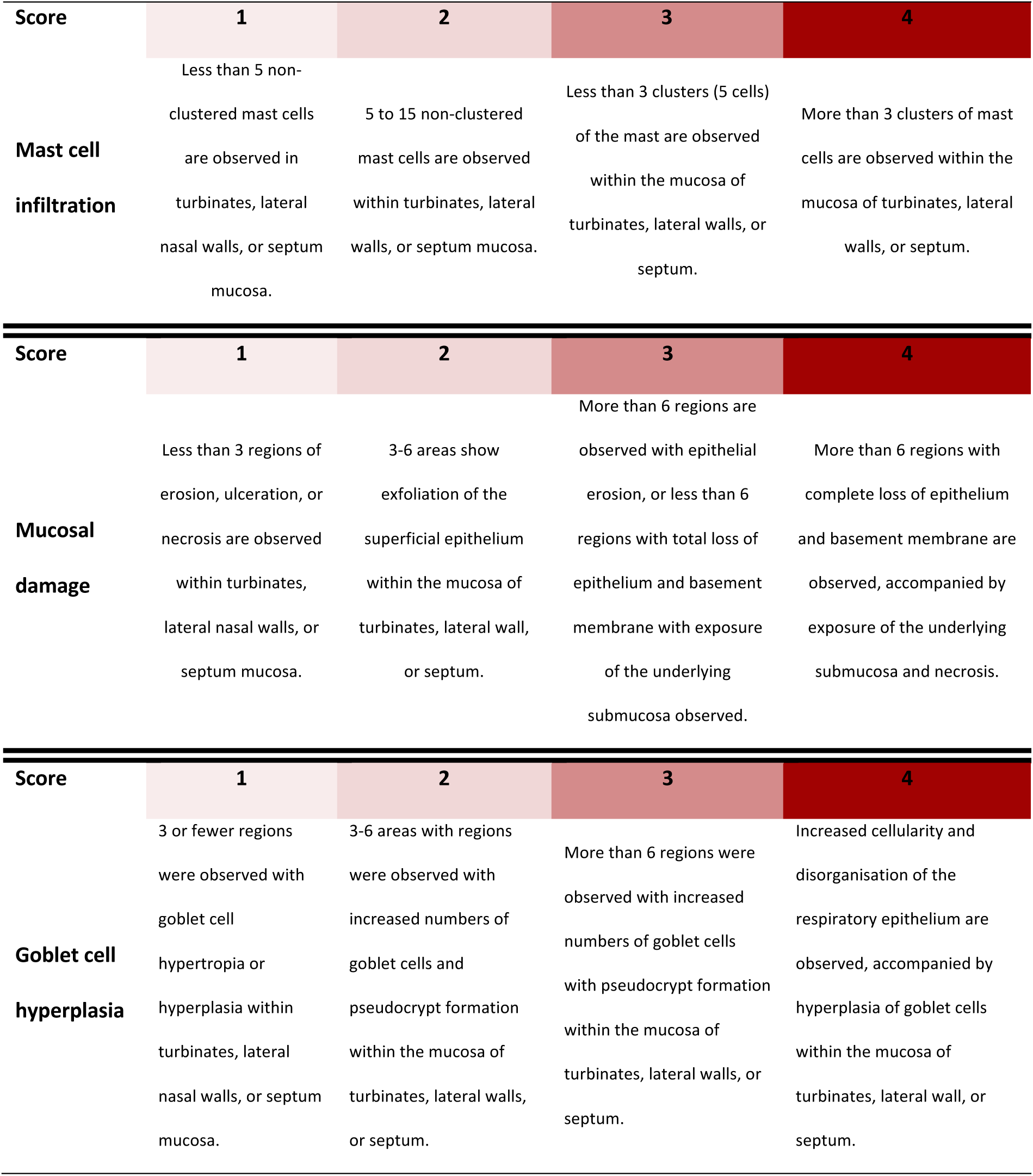
Semi-quantitative histopathologic score.

Eosinophilic and neutrophilic infiltration was quantified on H&E stained sections. In brief, 10 random regions of interest (ROI) with an area of 0.1 mm^2^ were created, overlaying the turbinates, lateral walls of the nasal cavity, or septum mucosa. The total number of cells within each ROI was quantified using cell segmentation based on nuclear detection (the hematoxylin channel). Cell segmentation was performed using the StarDist algorithm incorporated in Qupath (Weigert, Schmidt, Haase, Sugawara, & Myers, 2020). Eosinophils and neutrophils were manually annotated within each ROI.

T-lymphocytic infiltration was quantified in a semi-automated workflow on anti-CD3 stained sections. Ten random ROIs with an area of 0.2 mm^2^ were created, overlaying the turbinates, lateral borders of the nasal cavity, or septum mucosa. The total number of cells within the ROI was quantified based on nuclear detection (the DAPI channel) using the StarDist algorithm. Cells with Cy-5 signals above the set threshold (12500) were classified as CD3-positive T-lymphocytes. The samples were deidentified after the analysis to maintain experimental blinding of the groups.

### RNA extraction

The tissue samples stored in RNAlater Stabilisation Solution (Thermo Fisher Scientific, AM7020) were dissected, and the mucosal layer of the sinonasal cavity was collected for further processing. RNA was extracted from the mucosal tissue using the RNeasy kit (Qiagen, 74104, Hilden, Germany) according to the manufacturer’s instructions. The RNA integrity number (RIN) was analysed for each sample using the TapeStation System (Agilent, model 4200, Santa Clara, United States). The purity of RNA was assessed using the NanoDrop (Thermo Scientific Scientific, model ND-1000). The RNA concentration was quantified using the Qubit RNA High Sensitivity kit (Thermo Fisher Scientific, Q32852). After quality control, ribosomal RNA was removed from the extracted samples using the RiboMinus Eukaryote System v2 kit (Thermo Fisher Scientific, A15026) and concentrated with the RiboMinus Concentration Module kit (Thermo Fisher Scientific, K155005).

### Library preparation and sequencing

Long-read transcriptomics was performed on RNA extracts using the direct cDNA sequencing kit (Oxford Nanopore Technologies, SQK-DCS109, Oxford, UK) according to the manufacturer’s recommendations with the following minor modification: the protocol was started with 200 ng poly(A) enriched RNA. In short, after first-strand cDNAs were synthesised using the Maxima H Minus Reverse Transcriptase (Thermo Fisher Scientific, EP0752), the RNA was degraded with RNase Cocktail Enzyme Mix (Thermo Fisher Scientific, AM2286) and second-strand cDNAs were synthesised using the LongAmp Taq 2X Master Mix (New England Biolabs, M0287L, Ipswich, USA). The double-stranded cDNAs were end-repaired using the NEBNext Ultra II End Repair/dA-Tailing Module (New England Biolabs, E7546S). Sequencing adapters were ligated to the end-repaired strands using Blunt/TA Ligase Master Mix (New England Biolabs, M0367L). The cDNA libraries were purified after each enzymatic reaction using the AMPure XP beads (Beckman Coulter, A63882, Brea, US). All the samples were loaded on a SpotON flow cell (Oxford Nanopore Technologies, R9.4.1) and sequenced on a MinION (Oxford Nanopore Technologies, Mk1C).

### Transcriptomic quantification

FASTQ Base-calling was conducted with Guppy v 6.2.11 super accuracy mode using the ‘dna_r9.4.1_450bps_sup.cfg’ configuration (Oxford Nanopore Technology, Oxford UK). For each sample, all FASTQs were aggregated and run through a customised open-source Snakemake (Molder et al., 2021) and Snaketool (Roach et al., 2022) pipeline giantpandrna that can be accessed at https://github.com/gbouras13/giantpandrna. Input FASTQs were aligned with Minimap2 v 2.24 (Li, 2018) specifying ‘minimap2-ax splice’ to the Ensembl *Rattus norvegicus* release 108 top-level assembly and gtf file downloaded using the giantpandrna install command (Cunningham et al., 2022). The resulting bam files were sorted using Samtools v 1.16.1 (Li et al., 2009). These sorted bam files were input for transcriptome discovery and quantification using Bambu v3.0.0 (Chen et al., 2022).

### Immunoglobulin quantification

Immunoglobulin isotypes were mapped using a custom python program called NanoReceptor (https://github.com/gbouras13/NanoReceptor). Briefly, input transcripts were mapped to the IMGT database (Giudicelli et al., 2006) using Minimap2, filtered for mapped reads only using Samtools, and parsed to output Counts per Million transcript values for each Immunoglobulin isotype.

### Bioinformatics of transcriptomics

All subsequent analyses were performed in R v 4.2.0 (R Core Team, 2017). Differential gene expression analysis was conducted using the DESeq2 package v 1.38.3 from Bioconductor (Love, Huber, & Anders, 2014). The differentially expressed genes (DEGs) significance threshold was set at Benjamini–Hochberg adjusted p-value ≤0.05. Functions from the tidyverse collection of R packages v 1.3.2 were incorporated into the analysis and visualisation (Wickham et al., 2019). Pathway enrichment analysis was performed using the clusterProfiler 4.6.0 package (T. Wu et al., 2021). The functional enrichment analysis included the terms Gene Ontology (GO) (Mi, Muruganujan, Ebert, Huang, & Thomas, 2019), Kyoto Encyclopedia of Genes and Genomes (KEGG) (Kanehisa, Furumichi, Sato, Kawashima, & Ishiguro-Watanabe, 2023) and CellMarker 2.0 databases (Hu et al., 2023). For GO Over-representation analyses, all genes expressed in our dataset (n=12,878) were used as background. The GOSemSim R package v 2.24.0 was used to reduce redundancy among enriched GO terms, with a threshold of 0.7 (Yu et al., 2010). Only homologue ensemble genes of Homo sapiens and Rattus norvegicus with one-to-one orthologue correspondence were considered for the cell markers. The up and down-regulated genes for each cluster were separately analysed. All genes expressed in our dataset (n=10,384) were used as background. For the KEGG gene set enrichment (GSEA), pre-ranked GSEA was run on the list of genes, sorted by their log fold changes. The Benjamini-Hochberg method was used for p-value adjustment for all functional enrichment analyses, accounting for multiple testing. Only terms with a false discovery rate (FDR) ≤0.01 were considered significant.

### Bacterial genome sequencing

Whole-genome sequencing of the *S. aureus* clinical isolates was performed at a commercial sequencing facility (SA Pathology, Adelaide, SA, Australia) as previously described by Shaghayegh et al. (Shaghayegh et al., 2023). In short, the NextSeq 550 platform and the NextSeq 500/550 Mid-Output kit v2.5 (Illumina Inc. San Diego, USA) were used. Genomic DNA was isolated using the NucleoSpin Microbial DNA kit (Machery-Nagel GmbH and Co.KG, Duren, Germany). A modified protocol of the Nextera XT DNA library preparation kit (Illumina Inc.) was employed to develop sequencing libraries. Fragmentation of the genomic DNA and subsequent amplification of the Nextera XT indices to the DNA fragments were performed through a low-cycle PCR reaction. After manual purification and normalisation of the amplicon library, 150 bp reads were generated by sequencing. Using ABRicate (Seemann, Abricate), version 1.0.1, all isolate contigs were screened via Virulence Factor Database (Liu, Zheng, Jin, Chen, & Yang, 2019) to detect virulence genes.

### Data availability

The transcriptomics dataset from long-read sequencing is publicly accessible in the Sequence Read Archive (SRA) under accession number PRJNA910244.

### Statistics

Statistical analysis was performed in R v4.2.1 (R Core Team, 2017), with data expressed as mean ± standard error of the mean (s.e.m). The histopathological data were compared between groups using a one-way ANOVA test with a post-hoc t-test for lymphocyte and eosinophil count and a Kruskal-Wallis test with a post-hoc Dunnet test for the semi-quantitative scoring. The statistical significance threshold was set at p≤0.05, and p-values were corrected for multiple comparisons using the Benjamini-Hochberg method.

## Results

The *S. aureus* strains were isolated from CRS patients with high levels of CD45+ cells, 63% and 49% in the CRSwNP and CRSsNP patients, respectively. In contrast, the non-CRS carrier had a low count of CD45+ cells, indicating a non-inflamed mucosal tissue. Eosinophils were the dominant CD45+ cells in the CRSwNP patient (55%), while neutrophils comprised 65% of CD45+ cells in the CRSsNP patient (Figure S1B). These findings suggest a T2/eosinophilic endotype for the CRSwNP patient and a non-T2 endotype for the CRSsNP patient.

All rats in the 4 experimental groups survived the 28-day intranasal challenge with SABSFs without adverse events. The rats showed steady weight gain with increasing age, and no decrease in mean body weight was detected across the groups during the experiments or at the endpoint (data not shown).

### Long-term SABSF challenges induce multifocal inflammation

In order to assess the inflammation of the nasal mucosa after exposure to disease-specific *S. aureus* strains, we investigated the T-cell infiltration in the mucosa (epithelium and lamina propria) of the septum and turbinates. As shown in Figure 2A, T-cells were scattered within the epithelium, the lamina propria and the submucosa of the turbinates and the septum. T-cells within the lamina propria and submucosa of the turbinates were often multifocally distributed (Figure 2C), with low T-cell infiltrated regions in between (Figure 2B). The mean percentage of CD3-positive cells out of total cells in the septum and turbinates was 0.64 (± 0.05) for the vehicle control group. The CRSwNP strain group had the highest mean percentage of CD3-positive cells at 6.62% (± 0.8). A one-way ANOVA revealed a statistically significant difference between the groups. As shown in Figure 2D, compared to the vehicle control group, T-cell infiltration of the mucosa was significantly increased in all SABSF-treated groups (CRSwNP strain p≤0.01, CRSsNP strain p≤0.05, carrier strain p≤0.05).

**Figure 2.**
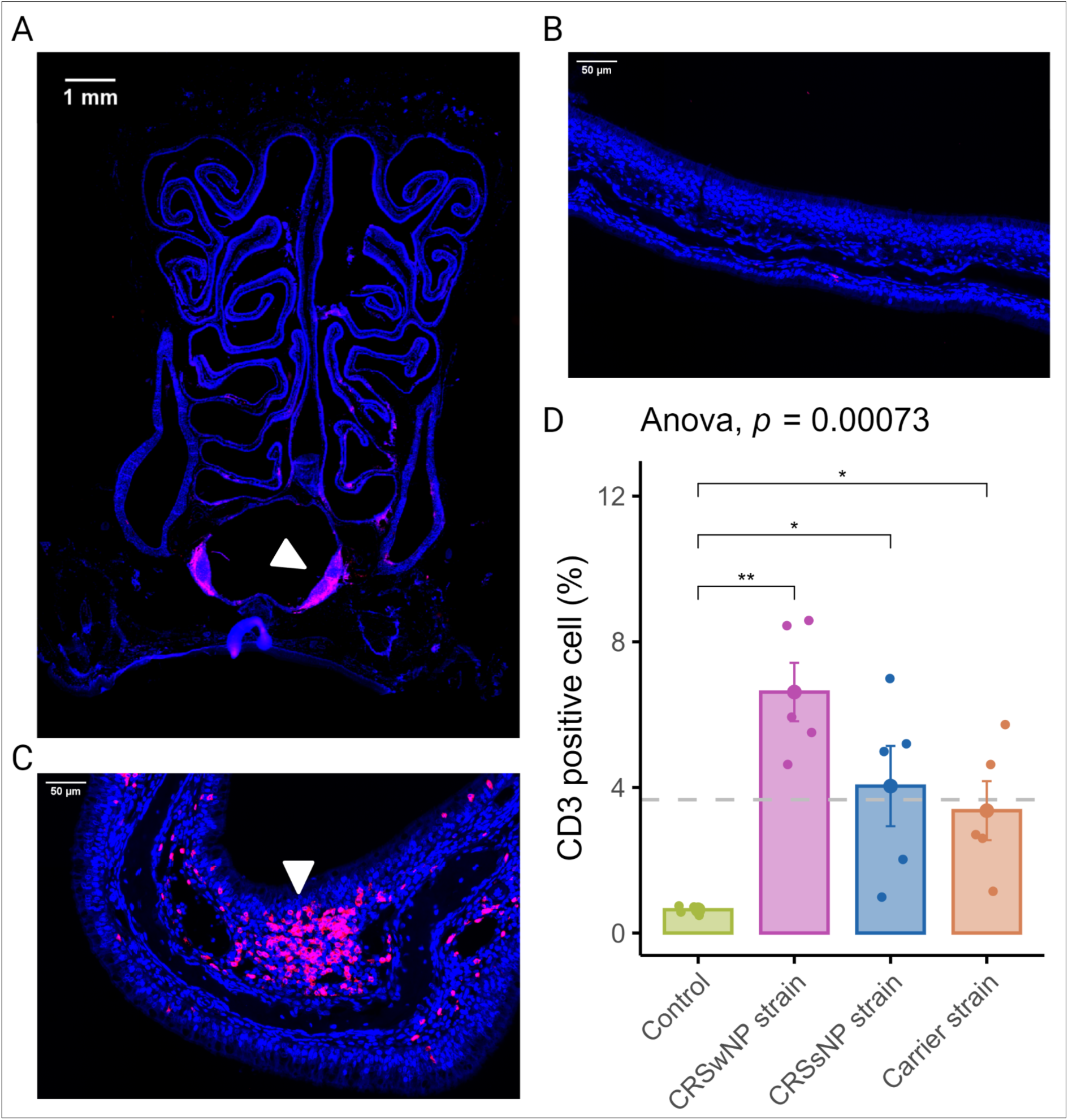
T-lymphocyte infiltration of the nasal mucosa after daily exposure to *S. aureus* biofilm-secreted factors (SABSF) (vehicle-control, SABSF CRSwNP strain, SABSF CRSsNP strain, or SABSF non-CRS carrier strain). (A) Representative immunofluorescent whole slide image of sinonasal coronal sections of a rat’s nose after daily challenge with SABSF with fluorescent detection of CD3 positive cells (pink, T-cell marker) and 4ʹ,6-diamidino-2-phenylindole (blue, DAPI; nuclear marker). The white arrow indicates the nasal-associated lymphoid tissue (NALT). Scale bar: 1mm. (B) Representative image of the nasal turbinate region with low T-cell infiltration of the mucosa. Scale bar: 50 µm. (C) Representative image of the nasal turbinate region with high T-cell infiltration of the mucosa (white arrow). Scale bar: 50 µm. (D) Percentage of T-cell counts per total cells in mucosa per group. The dashed line represents the mean of all samples. The points represent the individual measurements of the samples (mean of 10 ROI for n=5 per group). Error bars indicate mean±s.e.m. *p≤0.05, **p≤0.01, one-way ANOVA and pairwise post-hoc T-test with Benjamini-Hochberg p-value adjustment.

### Long-term SABSF challenges of nasal mucosa induce eosinophilic infiltration

CRSwNP predominantly has a T2 inflammation with a predominantly eosinophilic infiltrate. Eosinophils were quantified in coronal sections to evaluate the potential of SABSF to induce tissue eosinophilia (Figure 3A). Similarly to T-cells, eosinophilic mucosal infiltration was mainly seen to be multifocally distributed (Figure 3B-C). The mean rate of eosinophils observed in the nasal mucosa of the vehicle control animals was 0.96/1000 cells (± 0.35). In the SABSF groups, the mean eosinophil rate was 6.23 (±1.98), 5.00 (±0.90), and 2.04 (±0.29) for the animals stimulated with SABSF from the CRSwNP strain, CRSsNP strain and carrier strain, respectively. Interestingly, the difference between the vehicle control and CRSsNP strain groups was significant (p≤0.05)(Figure 3D). Neutrophils were less frequently observed in the nasal mucosa than eosinophils across all samples (2.77/1000 vs 3.55/1000). No significant correlation was found between the mean neutrophilic rate of the vehicle control and the SABSF groups (ANOVA, p = 0.25).

**Figure 3.**
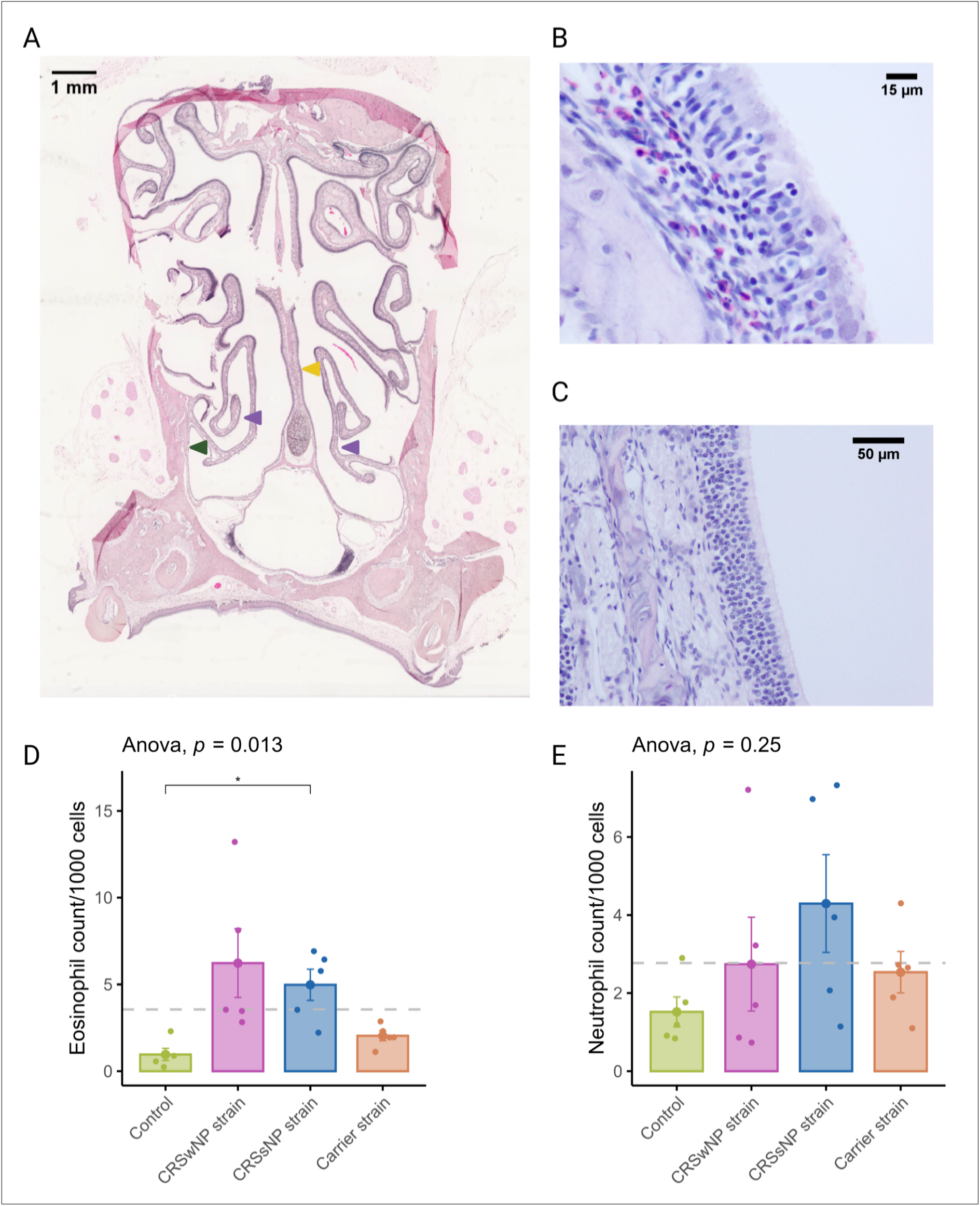
Eosinophil and neutrophil infiltration of the nasal mucosa after daily exposure to *S. aureus* biofilm-secreted factors (SABSF) (vehicle-control, SABSF CRSwNP strain, SABSF CRSsNP strain, or SABSF non-CRS carrier strain). (A) Representative histology whole slide image of sinonasal coronal sections of a rat after 28 days daily challenge with SABSF stained with H&E. Green arrow indicates lateral nasal wall, yellow arrow indicates nasal septum, and purple arrows indicate turbinates. Scale bar: 1 mm (B) Image of the nasal turbinate region showing a mixed inflammatory infiltrate in the lamina propria, predominantly composed of eosinophils and lymphocytes (the latter also infiltrating the lining epithelium), with fewer neutrophils. Scale bar: 15 µm. (C) Representative image of an unaffected area of the nasal turbinates with no inflammatory infiltrate present. Scale bar: 50 µm. (D) Quantification of the count (per 1000 cells) of eosinophils in mucosa per group. (E) Quantification of the count (per 1000 cells) of neutrophils in mucosa per group. The dashed line represents the mean of all samples. The points represent the individual measurements of the samples (mean of 10 ROI). Error bars indicate mean±s.e.m. Statistical analysis was performed by comparing groups to control using a one-way ANOVA and pairwise post-hoc T-test with Benjamini-Hochberg p-value adjustment (n=5 per group). (*p≤0.05).

### Long-term SABSF challenges induce mucosal damage and goblet cell hyperplasia of the nasal mucosa

It is now well-established that epithelial damage and barrier dysfunction are involved in the pathophysiology of CRS, and several mechanisms can play a role. One of the primary mechanisms leading to mucosal damage is the local accumulation of IgG, which induces the activation of the classical complement pathway and neutrophils (Kato, Schleimer, & Bleier, 2022). Another characteristic epithelial change observed in CRS is the increased expression of goblet or secretory cells (Burgel et al., 2000; Cho, Kim, & Gevaert, 2016). A semi-quantitative scoring system was used to evaluate the mucosal changes in the SABSF-challenged animals (Table 1). After 28 days of daily SABSF intranasal instillation, histopathology examination revealed mucosal damage for all 3 groups with focal epithelial erosion or ulceration (Figure 4A). Although some epithelial erosion was observed in the vehicle control group, most of the mucosa was intact. The mucosal damage in the CRSwNP strain group was significantly more compared to the vehicle control group (2.6, ±0.24 vs 1.4, ±0.24 in CRSwNP SABSF treated animals, and vehicle control treated animals, respectively, p≤0.05) (Figure 4B). Furthermore, histopathology revealed regions with proliferation and disorganisation of the respiratory epithelium combined with evidence of goblet cell hyperplasia in all 3 SABSF groups (Figure 4E). Goblet cell hyperplasia was significantly more in the CRSwNP strain group compared to the vehicle control group (3, ±0.45 vs 1, ±0, p≤0.05 in CRSwNP SABSF treated animals and vehicle control treated animals, respectively) (Figure 4F).

**Figure 4.**
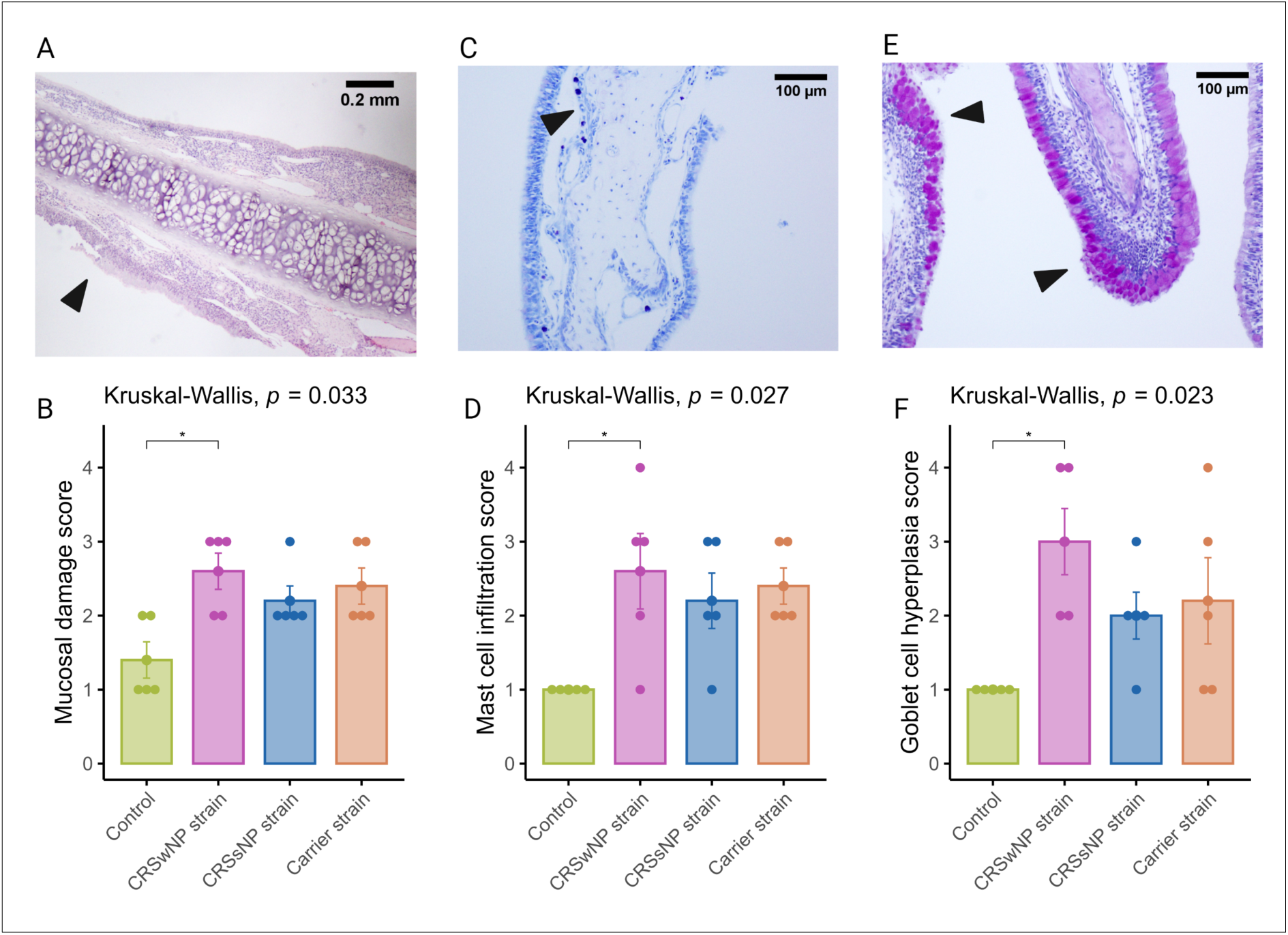
Semi-quantitative histopathologic score assessing mast cell infiltration, mucosal damage, and goblet cell hyperplasia after chronic exposure to *S. aureus* biofilm-secreted factors (SABSF) (vehicle-control, SABSF CRSwNP strain, SABSF CRSsNP strain, or SABSF non-CRS carrier strain) for 28 days (scoring scale, table 1). Mucosal damage was assessed in sections stained with H&E, mast cell infiltration in sections stained with toluidine blue, and goblet hyperplasia in sections with PAS staining. (A) Image of mucosal damage, black arrow indicates mucosal damage with loss of epithelium and basement membrane. Scale bar: 0.2 mm. (B) The mean score of mucosal damage per group. (C) Goblet cell hyperplasia (arrows) with proliferation and disorganisation of the respiratory epithelium. Scale bar: 100 µm. (C) The mean score of mast cell infiltration per group. (E) Image of goblet cell hyperplasia, the black arrows indicate proliferation and disorganisation of the respiratory epithelium with hyperplasia of goblet cells. Scale bar: 100 µm. (F) The mean score of goblet cell hyperplasia per group. The points represent the individual measurements of the samples. Error bars indicate mean±s.e.m. Statistical analysis was performed by comparing groups to control using a Kruskal-Wallis test and pairwise post-hoc Dunnet test with Benjamini-Hochberg p-value adjustment (n=5 per group) (*p≤0.05).

### Long-term SABSF challenges induce mastocytosis in the nasal mucosa

The current body of research examining the function of mast cells in CRS suggests that they may contribute to the pathophysiology of eosinophilic CRS, as evidenced by the significant increase in membrane-attached IgE-positive mast cells in patients with eosinophilic CRS compared to those with non-eosinophilic CRS (Baba, Kondo, Suzukawa, Ohta, & Yamasoba, 2017). Toluidine blue staining revealed aggregated mast cells in the nasal mucosa of SABSF-challenged rats, specifically in regions with mucosal damage (Figure 4C). Quantification of mast cells was significantly increased in the CRSwNP strain group compared to the vehicle control group (2.6, ±0.5 vs 0, ±0 in CRSwNP SABSF treated animals, and vehicle control treated animals, respectively, p≤0.05) (Figure 4F).

### Transcriptome profiling by RNA-seq reveals differential inflammation between SABSF groups

To comprehensively study the cellular transcriptional response of the nasal mucosa after SABSF challenges, long-read bulk RNA transcriptomics was performed. The transcriptional response was measured across a total of four conditions and three biological replicates. All comparisons were made to the vehicle control group. A total of 11,258,208; 11,463,508; 7,331,347; and 5,816,055 raw reads were obtained from the control, CRSwNP strain, CRSsNP strain, and carrier strain treated animals, respectively. Furthermore, 8,971,400; 9,635,611; 6,580,694; and 5,090,261 reads were mapped to the reference genome, respectively (Table S1). PCA separated the vehicle control group and SABSF groups on transcript level (1^st^ component 33.8% of the variance, 2^nd^ component 17.9% of the variance) and gene level (1^st^ component 39.9% of the variance, 2^nd^ component 17.4% of the variance). However, the expression profiles of the CRSsNP strain group samples were similar to those of the carrier strain group, suggesting a global similarity (Figure 5A). Differential expression analysis grouped by SABSF strain type revealed DEGs between all SABSF groups and vehicle control. The largest number of DEGs was found in the carrier strain group (n=449, adjusted p-value≤0.05) (Figure 5E), closely followed by the CRSsNP strain group (n=461, adjusted p-value≤0.05)(Figure 5D). Fewer DEGs were found in the CRSwNP strain group (n=221, adjusted p-value≤0.05)(Figure 5C). A set of 63 overlapping DEGs were found across all SABSF groups. In particular, a considerable overlap of DEGs between the CRSsNP strain group and the carrier strain group was observed, suggesting a convergent response between these two groups (Figure 5B). Detailed information on significant DEGs is shown in supplementary table

**Figure 5.**
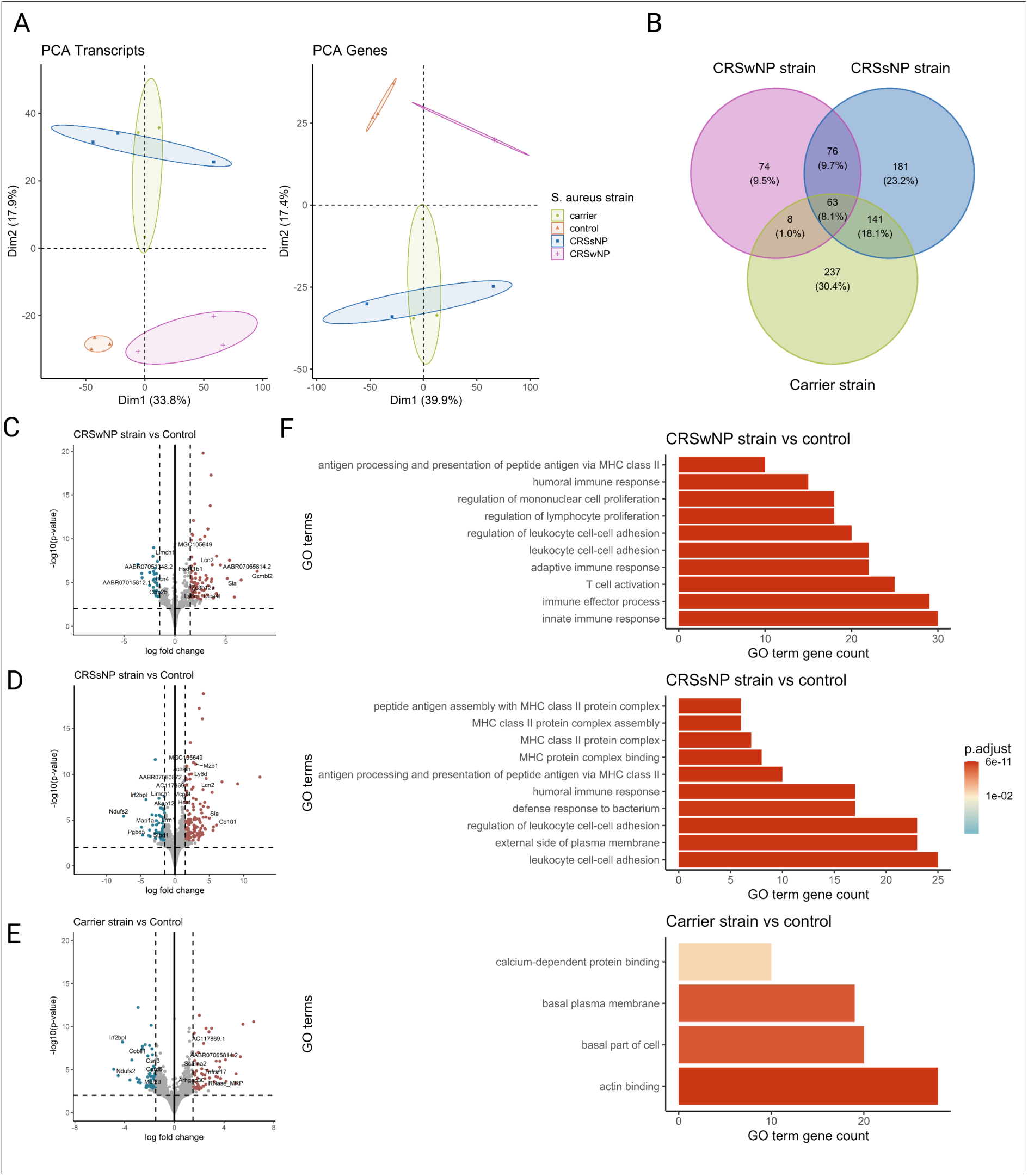
Transcriptome analysis of the nasal mucosal tissue from 12-week-old rats after chronic exposure to *S. aureus* biofilm-secreted factors (SABSF) for 28 days (vehicle-control (n=3), SABSF CRSwNP strain (n=3), SABSF CRSsNP strain (n=3), or SABSF non-CRS carrier strain (n=3)). (A) Principal component analysis based on the top 1000 most variable genes of nasal mucosa transcriptome data on transcript level (left) and gene level (right). Each point represents a sample and is coloured according to group. The numbers in parentheses on the x and y axis indicate the percentage of variation in the dataset explained by a principal component dimension (PC1 vs PC2). (B) Venn diagram shows the count with the percentage of the total between parentheses of unique differentially expressed genes (DEGs) and overlapping DEGs (adjusted p-value≤0.05) between the four groups. (C, D, E) Volcano plots of gene expression of all SABSF groups compared to control, significantly upregulated DEGs (adjusted p≤0.05) are indicated in red, and significantly downregulated DEGs (adjusted p≤0.05) are indicated in blue. Grey dots represent non-DEGs. Data are expressed as Log2 fold change. Vertical dashed lines are set at -1.5 and 1.5. The horizontal dashed line is set at 2. (F) Over-representation analysis of Gene Ontology (GO) pathways of DEGs (FDR p-value≤0.01). SABSF groups are compared to the control. Only the top 10 most significant pathways are shown per group. PCA, principal component analysis.

### Error! Reference source not found

Over-representation analysis was performed using GO pathway annotations to identify biological processes, molecular function and cell compartment genes enriched in the nasal mucosa of SABSF-challenged rats. A total of 99, 39, and 4 GO pathways were enriched for the CRSwNP, CRSsNP, and carrier strain group, respectively, compared to vehicle control. Despite DEGs differences, the over-representation analysis showed strikingly similar enriched pathways for the CRSwNP and CRSsNP strain groups. Among the DEGs, in both the CRSwNP strain group and CRSsNP group, we found enrichment of genes associated with immune effector process, innate immune response, antigen processing and presentation of exogenous antigen, leukocyte cell-cell adhesion, regulation of lymphocyte proliferation, T cell activation, humoral immune response, adaptive immune response, regulation of leukocyte cell-cell adhesion, pointing towards a similar biological process in the CRSwNP strain and CRSsNP strain challenged group. In contrast, no enriched pathways were implicated in immunity among the DEGs in response to SABSF challenges in the carrier strain group (Figure 5F). Detailed information on GO-enriched pathways is shown in supplementary table 3.

Next, to detect whole transcriptome expression perturbations comparing vehicle-control and SABSF-challenged groups, gene set enrichment analysis (GSEA) was performed. We used the KEGG gene sets to determine whether similarly coordinated changes to gene expression were observed along any of the pathways. Nine of the 30 significantly upregulated pathways were found in all SABSF groups (Table S4). Interestingly, the pathways terms are related to infection and immunity, such as “Staphylococcus aureus infection”, “Intestinal immune network for IgA production”, “Antigen processing and presentation”, and “Hematopoietic cell lineage”, indicating that immune response pathways are activated. However, The CRSwNP and CRSsNP strain groups included more pathways involved in infection and immunity, such as “Th1 and Th2 cell differentiation”, “Th17 cell differentiation”, “Allograft rejection”, and “Phagosome”, suggesting a more severe immune response (Figure 6A).

**Figure 6.**
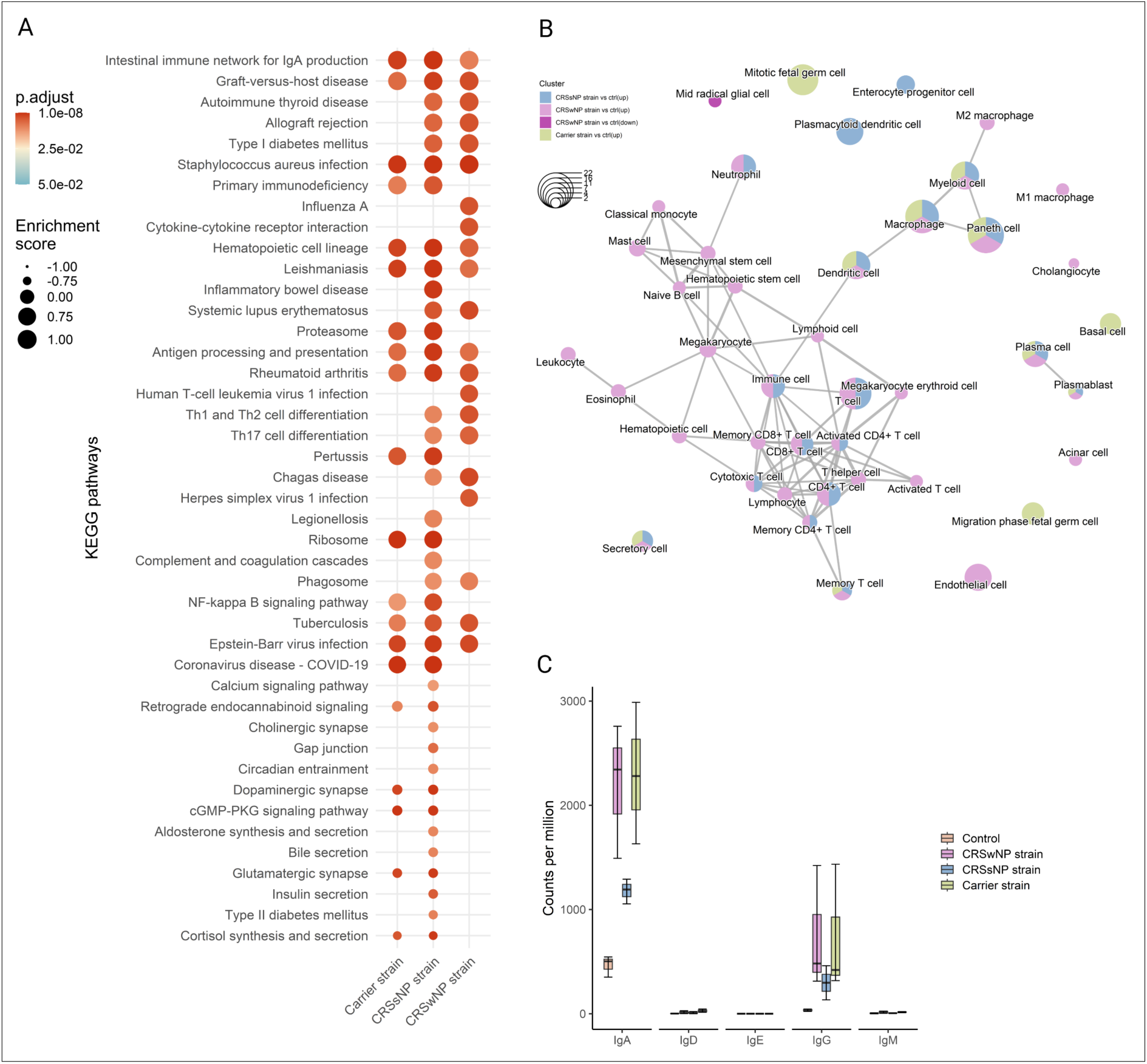
Functional gene expression profiles analysis of the nasal mucosal tissue from 12-week-old rats after chronic exposure to *S. aureus* biofilm-secreted factors (SABSF) for 28 days (vehicle-control (n=3), SABSF CRSwNP strain (n=3), SABSF CRSsNP strain (n=3), or SABSF non-CRS carrier strain (n=3)). (A) GSEA of pathways in SABSF-challenged nasal mucosa compared to vehicle controls using KEGG annotations. Only gene sets significantly altered (FDR-adjusted p≤0.01) compared to the vehicle control group are shown. Columns are ordered by enrichment score. (B) Enrichment network map of over-represented cell marker gene sets performed separately for upregulated and downregulated DEGs. Line width and the distance between connected gene sets indicate the similarity of gene sets. Node colour indicates the groups with the significantly enriched term, and node size indicates the count of DEGs count in the gene set. Only cell marker gene sets significantly enriched are shown (FDR-adjusted p≤0.01). (C) Transcript count (per million) of immunoglobulin heavy chain subtypes in the transcriptome of the nasal mucosal tissue after chronic exposure to *S. aureus* biofilm-secreted factors (SABSF). Natural killer T cell, NKT cell; Tumor-associated macrophage, TAM; Tissue-resident memory T cell, TRM cell; T follicular helper cell, Tfh cell; Innate lymphoid cell, ILC; Exhausted T cell, Tex cell; Exhausted CD4+ T cell, CD4+ Tex cell; Exhausted CD8+ T cell, CD8+ Tex cell.

Next, functional profile analysis for cell markers was performed to gain insights into the cell types underlying the biological processes of the nasal mucosa. Overwhelmingly, immune cell markers were enriched in upregulated DEGs of all SABSF-challenged groups (plasma cell, plasmablast, macrophage, dendritic cell, memory T cell) (Table S5). Notably, non-immune markers such as “secretory cell” were also enriched. Similarly to the KEGG functional analysis, enrichment results for the CRSwNP and CRSsNP strain groups were dominated by cell terms involved in immunity. Interestingly, the CRSwNP strain group had significantly enriched cell markers for eosinophils and mast cells (p≤0.05) (Figure 6B).

### Long-term SABSF challenges induce transcription of IgA and IgG

Finally, the antibody production in the nasal mucosa following SABSF challenges was investigated by mapping the IG heavy chain subtype transcripts. Clear segregation was observed between the four groups’ Ig transcripts. Most of the IG transcripts are coded for IgA or IgG subtypes. The CRSwNP and carrier strain groups had the most transcripts coding for IgA, followed by the CRSsNP group. The IgA subtype was predominantly expressed as IgA1 subclass antibodies (Figure S2). The IgG transcripts showed a similar divergence as IgA between the groups (Figure 6C). Interestingly, most IgG antibodies were expressed as the IgG2A subclass, except for the CRSsNP strain group (Figure S2). Notably, no transcripts were mapped to the IgE heavy chain.

### Virulence factors present in the bacterial genome

The virulence factors of *S. aureus* have been linked to the activation of different types of immune cells (Cheung, Bae, & Otto, 2021). We investigated these virulence factors in the bacterial genomes and found that the CRSsNP strain contained 54, the CRSwNP strain had 52, and the non-CRS carrier strain had 62 known virulence genes. Notably, the carrier strain was found to carry the superantigen TSST-1 (Figure S3).

## Discussion

This study compared the histopathological and transcriptomic response to intranasal SABSF challenges with *S. aureus* strains isolated from CRS (CRSwNP, CRSsNP) and a non-CRS carrier strain *in vivo* in a chronic rodent model. In chronic infections, *S. aureus* is often found in biofilm formations (Hall-Stoodley, Costerton, & Stoodley, 2004; H. Wu, Moser, Wang, Hoiby, & Song, 2015). Furthermore, biofilm presence seems to be an independent factor for disease severity, the persistence of postoperative symptoms, ongoing mucosal inflammation, and infections in CRS patients (Singhal, Psaltis, Foreman, & Wormald, 2010). Moreover, the expression of genes differs significantly between planktonic and biofilm modes of life for *S. aureus* (Resch, Rosenstein, Nerz, & Gotz, 2005), and it has been demonstrated that exoproteins isolated from *S. aureus* biofilm can induce inflammation and negatively affects the viability and mucosal barrier of nasal epithelial cell cultures (Malik et al., 2015; Panchatcharam et al., 2020). Furthermore, recent studies have shown that *in vitro S. aureus* biofilm properties, such as the quantity of exoproteins produced by those biofilms associated with levels of inflammation and the localisation of inflammatory cells in CRS patients (Shaghayegh et al., 2022). Therefore, the SABSF used in this study were isolated from mature 48-hour *S. aureus* biofilm grown *in vitro*. However, since planktonic cells are typically dispersed from such mature biofilms at this growth stage, the SABSF harvested and used in our experiments likely comprises some exoproteins secreted by *S. aureus* planktonic cells.

Histopathological analysis showed that chronic intranasal challenges with SABSF induced inflammation, evidenced by significant T-lymphocytic infiltration in the nasal mucosa of the turbinates for all CIs tested compared to vehicle control treated animals. Interestingly, the inflammation of the mucosal lining was multifocally rather than diffusely distributed. In accordance with the present results, previous studies have also demonstrated an increase of inflammatory cells, such as lymphocytes, neutrophils, eosinophils, and plasma cells, in the subepithelial layer and lamina propria of the nasal mucosa after chronic exposure to *S. aureus* (Boase, Valentine, Singhal, Tan, & Wormald, 2011; Jia et al., 2014). Interestingly, various inflammatory cell types aggregates have also been demonstrated in human nasal tissue and nasal polyps (Shaghayegh et al., 2022). This study significantly expanded eosinophils after chronic stimulation with SABSF isolated from a CRSsNP patient. The expansion of eosinophilic cells in animals challenged with the SABSF from the other two groups showed more variability between animals tested and did not reach statistical significance. Regardless, from the 3 CIs tested, the SABSF harvested from *S. aureus* isolated from a CRSwNP patient appeared to have the greatest inflammatory propensity, evidenced by the presence of significant mucosal damage, goblet cell hyperplasia, and mast cell infiltration compared to the control. Because SABSF challenges from the three strains were dosed equally, this indicates that SABSF from a *S. aureus* isolated from a CRSwNP patient contained qualitatively different or more factors that can account for this mucosal damage and inflammation. The pathogenicity of *S. aureus* strains is determined indeed by their ability to produce virulence factors. It was therefore somewhat surprising that most virulence factors genes were carried by the carrier strain genome, including Toxic shock syndrome toxin-1, which was not present in the other strains. A potential difference in virulence factors expression profile between isolates might explain this observation.

To further pursue the observed difference in inflammation after stimulation to SABSF, an unbiased approach was employed to investigate the gene expression variation of the inflammatory response of the nasal mucosa. GO, KEGG, and cell marker gene sets functional enrichment analysis of the mucosal transcriptome showed a consistent response to the different stimuli across the three SABSF groups, where the primary sources of variation can be attributed to the SABSF group type. The SABSF from a *S. aureus* isolated from a CRSwNP patient consistently included more infection and immunity pathways corroborating the previously discussed histopathological findings.

An initial objective of the project was to assess the production of IgE in the nasal mucosa after chronic exposure to SABSF, as several studies have shown the presence of *S. aureus*-specific IgE in CRS (Foreman et al., 2011; Gevaert et al., 2005; Takeda et al., 2019). The results of this study indicate that exposure to SABSF did not lead to the local production in the nasal mucosa of IgE. Instead, an increase was seen in IgA and IgG production. Ig class switching to IgE is regulated by IL-4, a type 2 immunity cytokine significantly increased in e-CRS (Junttila, 2018; Kato, Peters, et al., 2022). The absence of IgE production might be related to insufficient IL-4 secretion (Th2 cells), as SABSF nasal challenges did not induce a stark type 2 immune response. This also accords with the observations of IgG subtypes, which showed that the majority of the IgG was IgG2 as a type 1 immunity induces the production of IgG2a, whereas the type 2 immunity stimulates the expression of IgG1, rendering each isotype an indicator of the underlying immune response (Firacative et al., 2018). These findings suggest that prolonged *S. aureus* infection/exposure in the nasal cavity might not independently lead to Th cell polarisation and IgE production. It might be that the local *S. aureus*-specific IgE production in CRS occurs in type 2 inflammatory states of the nasal tissue preceding or co-occurring with a *S. aureus* infection, leading to the allergic sensitisation to *S. aureus* secreted factors. Further studies, which consider these variables, will need to be undertaken.

## Abbreviations

CRS: Chronic rhinosinusitis
CRSsNP: CRS without nasal polyps
CRSwNP: CRS with nasal polyps
DEG: differentially expressed genes
GSEA: gene set enrichment
H&E: hematoxylin and eosin
IHC-IF: immunohistochemistry staining using immunofluorescence detection
IL: interleukin
MFU: McFarland Units
NALT: nasal-associated lymphoid tissue
PAS: periodic acid-Schiff
Rcf: relative centrifugal force
RIN RNA: integrity number
ROI: regions of interest
s.e.m: standard error of the mean
SABSF: *S. aureus* biofilm-secreted factors
Th: T-helper
TSLP: Thymic stromal lymphopoietin
T2: type 2

---

**Figure S1.**
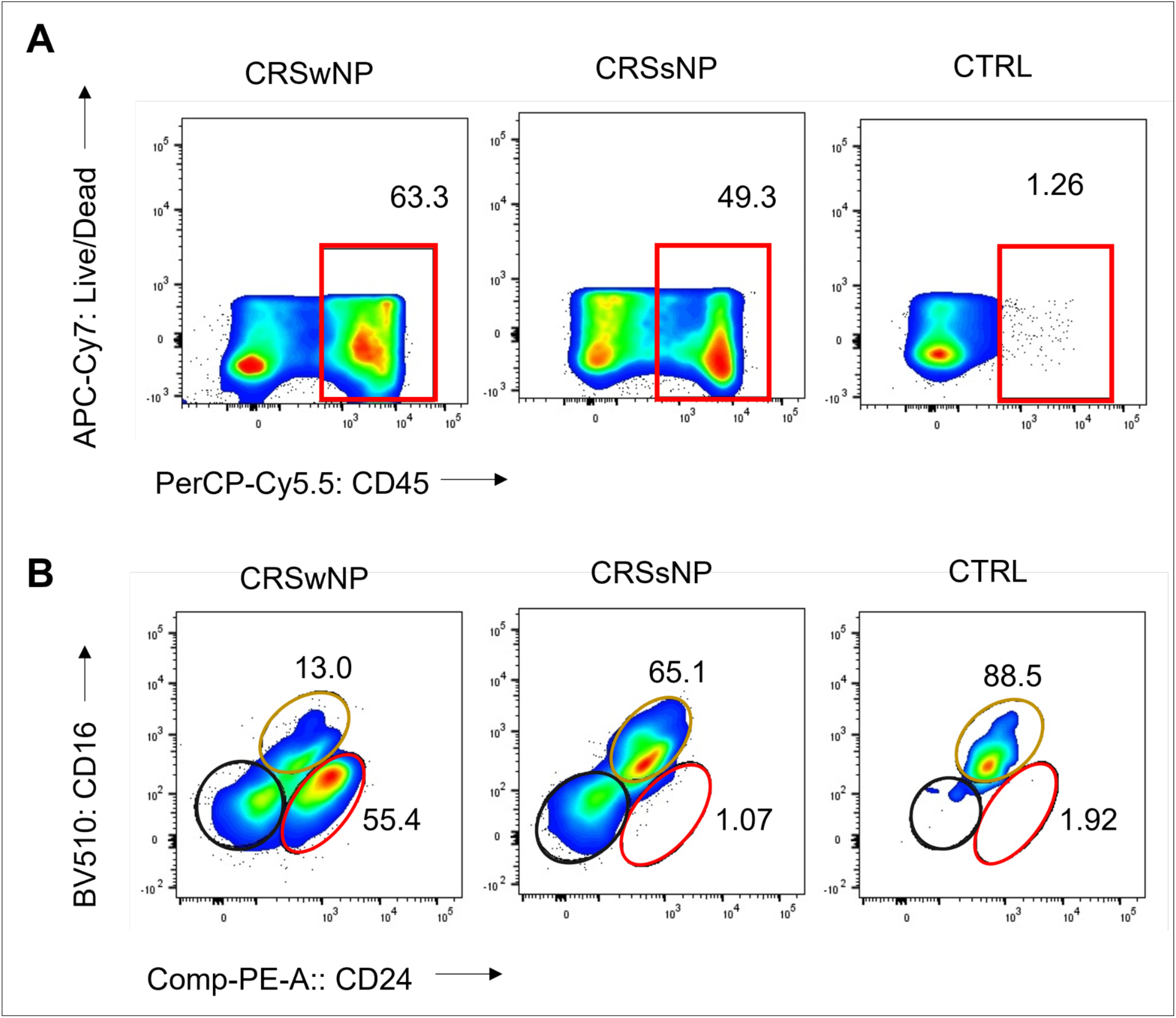
Frequency of CD45+ cells, eosinophils, and neutrophils in sinonasal tissue of CRS patients and non-CRS control. (A) Flow cytometry plots show frequency values and cell type gating. The gating of live leukocytes was performed using anti-CD45 PerCP-Cy5.5 antibody and Live/Dead APC-Cy7, following nucleated cell identification via FSC-A versus SSC-A plot and single-cell identification via FSC-A versus FSC-H plot. Eosinophils were identified as CD16-CD24+, neutrophils as CD16+ CD24-, and mast cells as CD16-CD24- from CD45+ and SSC-high granulocytes (Alexa Flour 488-CD14 and SSC-A). (B) The frequency of CD45+ cells, eosinophils (represented by the red line), neutrophils (represented by the yellow line), and mast cells (represented by the black line) is shown.

**Figure S2.**
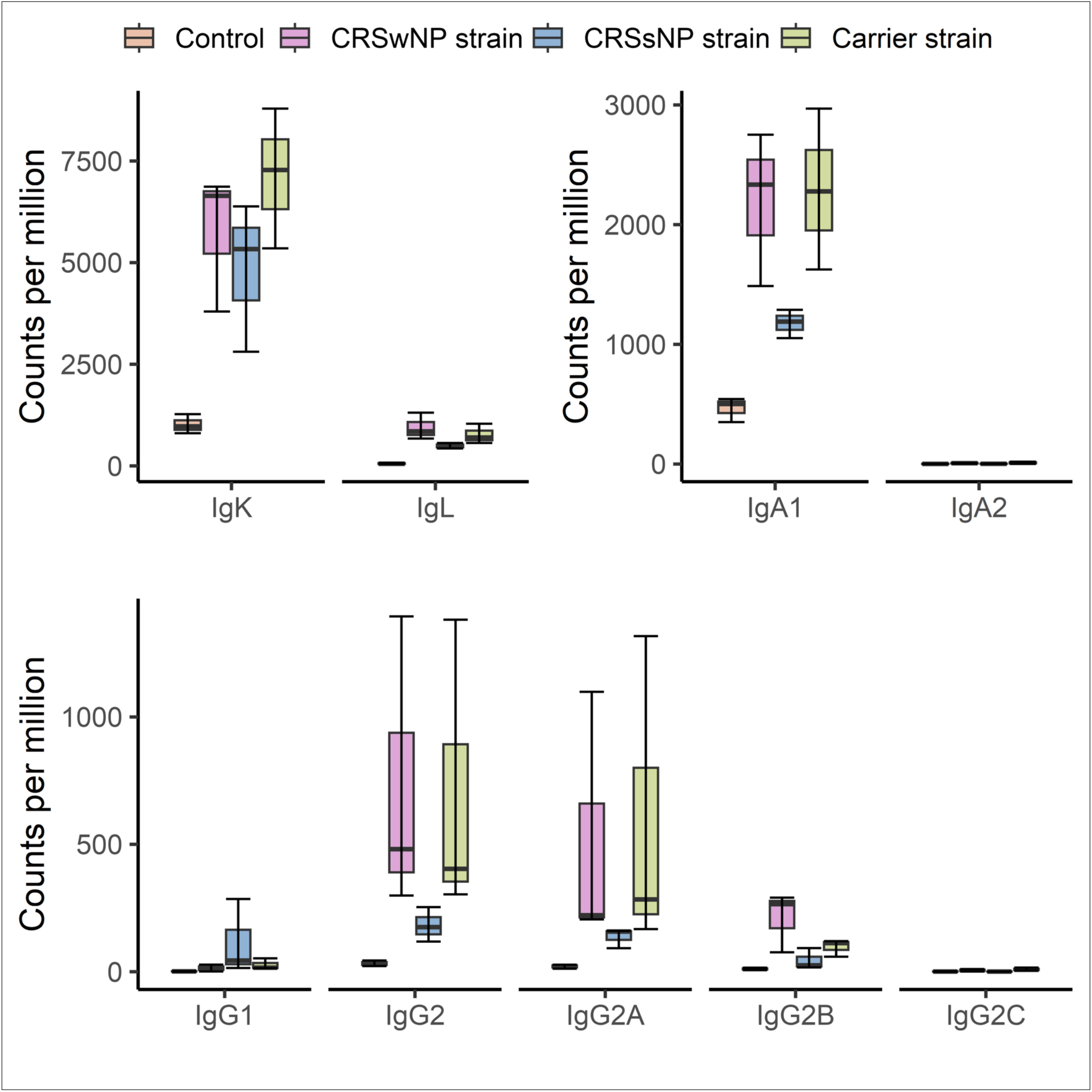
Nasal mucosal tissue transcriptome analysis revealed a differential expression of immunoglobulin light chain, IgA, and IgG subtypes after chronic exposure to *S. aureus* biofilm-secreted factors (SABSF). The transcript count (per million) of the subclasses is presented.

**Figure S3.**
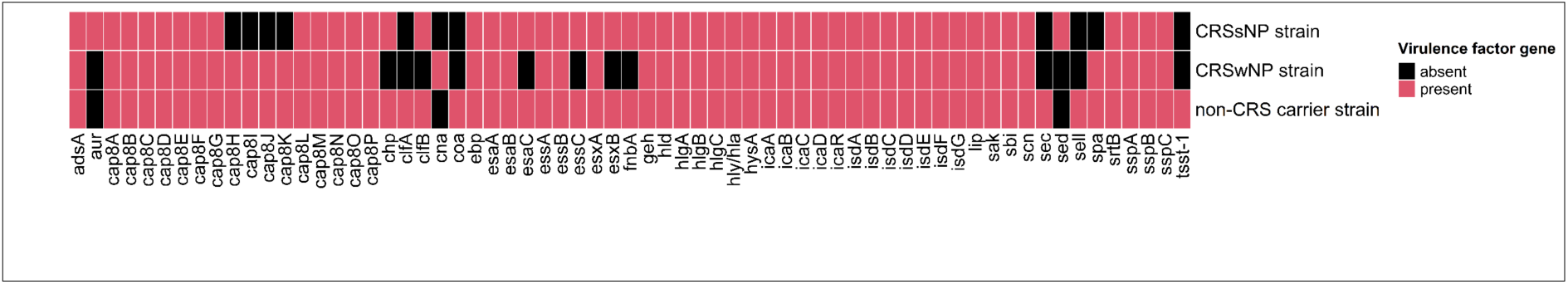
Matrix showing the presence of virulence genes within the genome of clinical isolates identified by Whole-Genome Sequencing and screening against the Virulence Factor Database (VFDB).

**Table S1.**
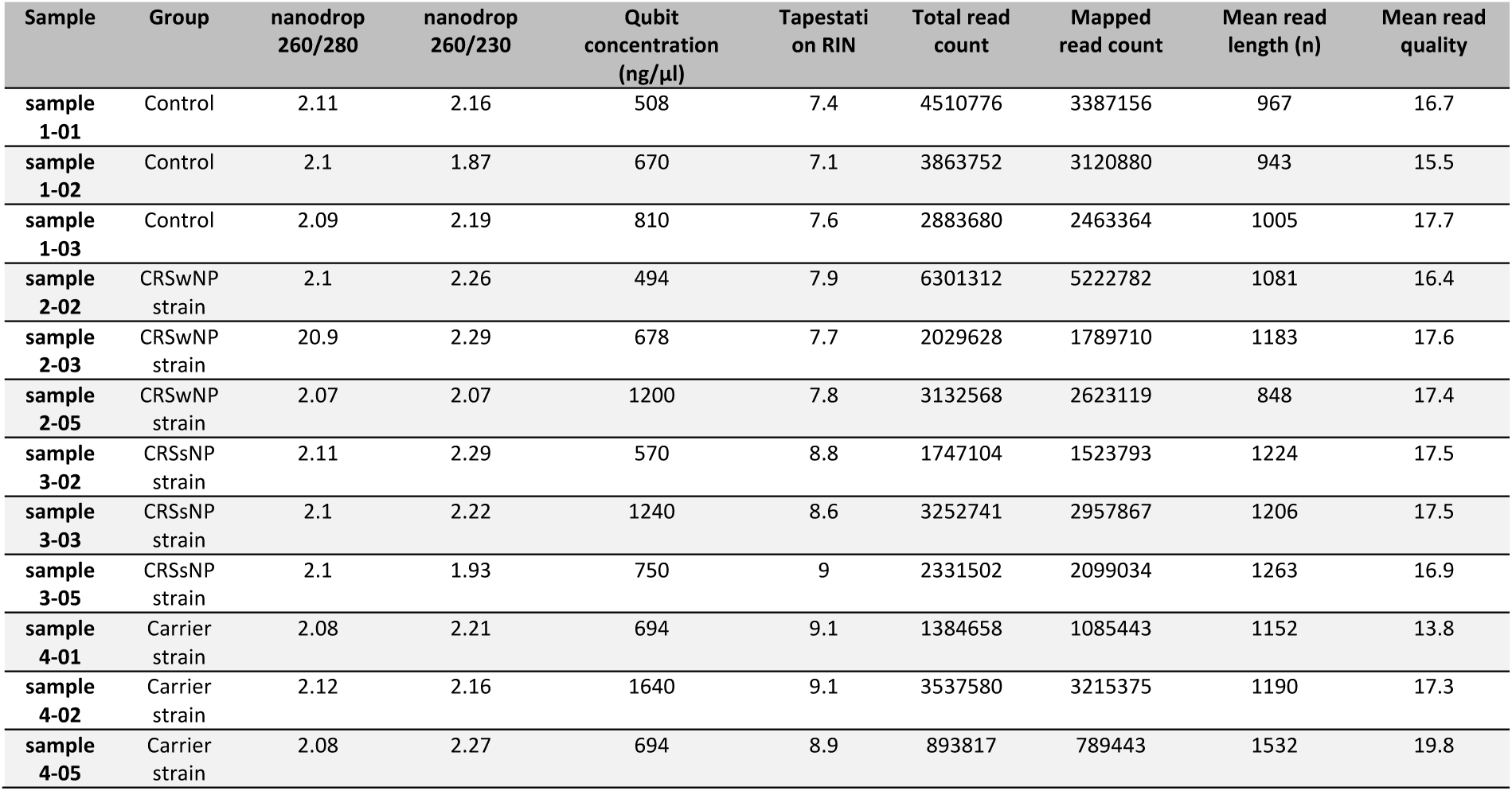
Quality control parameters transcriptomics.

**Table S2. Significant differently expressed genes**

Dataset is avialiabe on Figshare (an online open access repository) under the following link: https://figshare.com/s/bf773a2a29101334949f

**Table S3. Significant gene ontology pathways**

Dataset is avialiabe on Figshare (an online open access repository) under the following link: https://figshare.com/s/eb0b8bdc136aa2f61b54

**Table S4. Significant KEGG gene set enrichment pathways**

Dataset is avialiabe on Figshare (an online open access repository) under the following link: https://figshare.com/s/a0f35ce455fe99e91f89

**Table S5. Significant cell marker pathways**

Dataset is avialiabe on Figshare (an online open access repository) under the following link: https://figshare.com/s/babc21e7d369ac27c27f

## References

Baba, S., Kondo, K., Suzukawa, M., Ohta, K., & Yamasoba, T. (2017). Distribution, subtype population, and IgE positivity of mast cells in chronic rhinosinusitis with nasal polyps. Ann Allergy Asthma Immunol, 119(2), 120–128. doi:10.1016/j.anai.2017.05.019

Bachert, C., Gevaert, P., Holtappels, G., Johansson, S. G., & van Cauwenberge, P. (2001). Total and specific IgE in nasal polyps is related to local eosinophilic inflammation. J Allergy Clin Immunol, 107(4), 607–614. doi:10.1067/mai.2001.112374

Bankhead, P., Loughrey, M. B., Fernandez, J. A., Dombrowski, Y., McArt, D. G., Dunne, P. D.,… Hamilton, P. W. (2017). QuPath: Open source software for digital pathology image analysis. Sci Rep, 7(1), 16878. doi:10.1038/s41598-017-17204-5

Boase, S., Valentine, R., Singhal, D., Tan, L. W., & Wormald, P. J. (2011). A sheep model to investigate the role of fungal biofilms in sinusitis: fungal and bacterial synergy. Int Forum Allergy Rhinol, 1(5), 340–347. doi:10.1002/alr.20066

Burgel, P. R., Escudier, E., Coste, A., Dao-Pick, T., Ueki, I. F., Takeyama, K.,… Nadel, J. A. (2000). Relation of epidermal growth factor receptor expression to goblet cell hyperplasia in nasal polyps. J Allergy Clin Immunol, 106(4), 705–712. doi:10.1067/mai.2000.109823

Chen, Y., Sim, A., Wan, Y. K., Yeo, K., Lee, J. J. X., Ling, M. H.,… Göke, J. (2022). Context-Aware Transcript Quantification from Long Read RNA-Seq data with Bambu. bioRxiv, 2022.2011.2014.516358. doi:10.1101/2022.11.14.516358

Cheung, G. Y. C., Bae, J. S., & Otto, M. (2021). Pathogenicity and virulence of Staphylococcus aureus. Virulence, 12(1), 547–569. doi:10.1080/21505594.2021.1878688

Cho, S. H., Kim, D. W., & Gevaert, P. (2016). Chronic Rhinosinusitis without Nasal Polyps. J Allergy Clin Immunol Pract, 4(4), 575–582. doi:10.1016/j.jaip.2016.04.015

Cunningham, F., Allen, J. E., Allen, J., Alvarez-Jarreta, J., Amode, M. R., Armean, I. M.,… Flicek, P. (2022). Ensembl 2022. Nucleic Acids Res, 50(D1), D988–D995. doi:10.1093/nar/gkab1049

Firacative, C., Gressler, A. E., Schubert, K., Schulze, B., Muller, U., Brombacher, F.,… Alber, G. (2018). Identification of T helper (Th)1- and Th2-associated antigens of Cryptococcus neoformans in a murine model of pulmonary infection. Sci Rep, 8(1), 2681. doi:10.1038/s41598-018-21039-z

Fokkens, W. J., Lund, V. J., Hopkins, C., Hellings, P. W., Kern, R., Reitsma, S.,… Zwetsloot, C. P. (2020). European Position Paper on Rhinosinusitis and Nasal Polyps 2020. Rhinology, 58(Suppl S29), 1–464. doi:10.4193/Rhin20.600

Foreman, A., Holtappels, G., Psaltis, A. J., Jervis-Bardy, J., Field, J., Wormald, P. J., & Bachert, C. (2011). Adaptive immune responses in Staphylococcus aureus biofilm-associated chronic rhinosinusitis. Allergy, 66(11), 1449–1456. doi:10.1111/j.1398-9995.2011.02678.x

Gevaert, P., Holtappels, G., Johansson, S. G., Cuvelier, C., Cauwenberge, P., & Bachert, C. (2005). Organisation of secondary lymphoid tissue and local IgE formation to Staphylococcus aureus enterotoxins in nasal polyp tissue. Allergy, 60(1), 71–79. doi:10.1111/j.1398-9995.2004.00621.x

Gevaert, P., Nouri-Aria, K. T., Wu, H., Harper, C. E., Takhar, P., Fear, D. J.,… Gould, H. J. (2013). Local receptor revision and class switching to IgE in chronic rhinosinusitis with nasal polyps. Allergy, 68(1), 55–63. doi:10.1111/all.12054

Giudicelli, V., Duroux, P., Ginestoux, C., Folch, G., Jabado-Michaloud, J., Chaume, D., & Lefranc, M. P. (2006). IMGT/LIGM-DB, the IMGT comprehensive database of immunoglobulin and T cell receptor nucleotide sequences. Nucleic Acids Res, 34(Database issue), D781–784. doi:10.1093/nar/gkj088

Grayson, J. W., Cavada, M., & Harvey, R. J. (2019). Clinically relevant phenotypes in chronic rhinosinusitis. J Otolaryngol Head Neck Surg, 48(1), 23. doi:10.1186/s40463-019-0350-y

Hall-Stoodley, L., Costerton, J. W., & Stoodley, P. (2004). Bacterial biofilms: from the natural environment to infectious diseases. Nat Rev Microbiol, 2(2), 95–108. doi:10.1038/nrmicro821

Hastan, D., Fokkens, W. J., Bachert, C., Newson, R. B., Bislimovska, J., Bockelbrink, A.,… Burney, P. (2011). Chronic rhinosinusitis in Europe - an underestimated disease. A GA2LEN study. Allergy, 66(9), 1216–1223. doi:10.1111/j.1398-9995.2011.02646.x

Herbert, R. A., Janardhan, K. S., Pandiri, A. R., Cesta, M. F., & Miller, R. A. (2018). Nose, Larynx, and Trachea. In A. W. Suttie (Ed.), Boorman’s Pathology of the Rat (pp. 391–435). Boston: Academic Press.

Hu, C., Li, T., Xu, Y., Zhang, X., Li, F., Bai, J.,… Zhang, Y. (2023). CellMarker 2.0: an updated database of manually curated cell markers in human/mouse and web tools based on scRNA-seq data. Nucleic Acids Res, 51(D1), D870–D876. doi:10.1093/nar/gkac947

Hubrecht, R. (2013). Revised Australian code for the care and use of animals for scientific purposes. Animal welfare, 22(4), 491–491. doi:10.1017/S0962728600005674

Jia, M., Chen, Z., Du, X., Guo, Y., Sun, T., & Zhao, X. (2014). A simple animal model of Staphylococcus aureus biofilm in sinusitis. Am J Rhinol Allergy, 28(2), e115–119. doi:10.2500/ajra.2014.28.4030

Junttila, I. S. (2018). Tuning the Cytokine Responses: An Update on Interleukin (IL)-4 and IL-13 Receptor Complexes. Front Immunol, 9, 888. doi:10.3389/fimmu.2018.00888

Kanehisa, M., Furumichi, M., Sato, Y., Kawashima, M., & Ishiguro-Watanabe, M. (2023). KEGG for taxonomy-based analysis of pathways and genomes. Nucleic Acids Res, 51(D1), D587–D592. doi:10.1093/nar/gkac963

Kato, A., Peters, A. T., Stevens, W. W., Schleimer, R. P., Tan, B. K., & Kern, R. C. (2022). Endotypes of chronic rhinosinusitis: Relationships to disease phenotypes, pathogenesis, clinical findings, and treatment approaches. Allergy, 77(3), 812–826. doi:10.1111/all.15074

Kato, A., Schleimer, R. P., & Bleier, B. S. (2022). Mechanisms and pathogenesis of chronic rhinosinusitis. J Allergy Clin Immunol, 149(5), 1491–1503. doi:10.1016/j.jaci.2022.02.016

Lan, F., Zhang, N., Holtappels, G., De Ruyck, N., Krysko, O., Van Crombruggen, K.,… Bachert, C. (2018). Staphylococcus aureus Induces a Mucosal Type 2 Immune Response via Epithelial Cell-derived Cytokines. Am J Respir Crit Care Med, 198(4), 452–463. doi:10.1164/rccm.201710-2112OC

Li, H. (2018). Minimap2: pairwise alignment for nucleotide sequences. Bioinformatics, 34(18), 3094–3100. doi:10.1093/bioinformatics/bty191

Li, H., Handsaker, B., Wysoker, A., Fennell, T., Ruan, J., Homer, N.,… Genome Project Data Processing, S. (2009). The Sequence Alignment/Map format and SAMtools. Bioinformatics, 25(16), 2078–2079. doi:10.1093/bioinformatics/btp352

Liu, B., Zheng, D., Jin, Q., Chen, L., & Yang, J. (2019). VFDB 2019: a comparative pathogenomic platform with an interactive web interface. Nucleic Acids Res, 47(D1), D687–D692. doi:10.1093/nar/gky1080

Love, M. I., Huber, W., & Anders, S. (2014). Moderated estimation of fold change and dispersion for RNA-seq data with DESeq2. Genome Biol, 15(12), 550. doi:10.1186/s13059-014-0550-8

Malik, Z., Roscioli, E., Murphy, J., Ou, J., Bassiouni, A., Wormald, P. J., & Vreugde, S. (2015). Staphylococcus aureus impairs the airway epithelial barrier *in vitro*. Int Forum Allergy Rhinol, 5(6), 551–556. doi:10.1002/alr.21517

Mi, H., Muruganujan, A., Ebert, D., Huang, X., & Thomas, P. D. (2019). PANTHER version 14: more genomes, a new PANTHER GO-slim and improvements in enrichment analysis tools. Nucleic Acids Res, 47(D1), D419–D426. doi:10.1093/nar/gky1038

Miljkovic, D., Bassiouni, A., Cooksley, C., Ou, J., Hauben, E., Wormald, P. J., & Vreugde, S. (2014). Association between group 2 innate lymphoid cells enrichment, nasal polyps and allergy in chronic rhinosinusitis. Allergy, 69(9), 1154–1161. doi:10.1111/all.12440

Molder, F., Jablonski, K. P., Letcher, B., Hall, M. B., Tomkins-Tinch, C. H., Sochat, V.,… Koster, J. (2021). Sustainable data analysis with Snakemake. F1000Res, 10, 33. doi:10.12688/f1000research.29032.2

Mulcahy, M. E., Leech, J. M., Renauld, J. C., Mills, K. H., & McLoughlin, R. M. (2016). Interleukin-22 regulates antimicrobial peptide expression and keratinocyte differentiation to control Staphylococcus aureus colonisation of the nasal mucosa. Mucosal Immunol, 9(6), 1429–1441. doi:10.1038/mi.2016.24

Okifo, O., Ray, A., & Gudis, D. A. (2022). The Microbiology of Acute Exacerbations in Chronic Rhinosinusitis - A Systematic Review. Front Cell Infect Microbiol, 12, 858196. doi:10.3389/fcimb.2022.858196

Panchatcharam, B. S., Cooksley, C. M., Ramezanpour, M., Vediappan, R. S., Bassiouni, A., Wormald, P. J.,… Vreugde, S. (2020). Staphylococcus aureus biofilm exoproteins are cytotoxic to human nasal epithelial barrier in chronic rhinosinusitis. Int Forum Allergy Rhinol, 10(7), 871–883. doi:10.1002/alr.22566

R Core Team. (2017). R: A language and environment for statistical computing. R Foundation for Statistical Computing, Vienna, Austria.

Resch, A., Rosenstein, R., Nerz, C., & Gotz, F. (2005). Differential gene expression profiling of Staphylococcus aureus cultivated under biofilm and planktonic conditions. Appl Environ Microbiol, 71(5), 2663–2676. doi:10.1128/AEM.71.5.2663-2676.2005

Roach, M. J., Pierce-Ward, N. T., Suchecki, R., Mallawaarachchi, V., Papudeshi, B., Handley, S. A.,… Edwards, R. A. (2022). Ten simple rules and a template for creating workflows-as-applications. PLOS Computational Biology, 18(12), e1010705. doi:10.1371/journal.pcbi.1010705

Seemann, T. (Abricate). Abricate. Retrieved from https://github.com/tseemann/abricate

Shaghayegh, G., Cooksley, C., Bouras, G. S., Panchatcharam, B. S., Idrizi, R., Jana, M.,… Vreugde, S. (2022). Chronic rhinosinusitis patients display an aberrant immune cell localisation with enhanced S aureus biofilm metabolic activity and biomass. Journal of Allergy and Clinical Immunology. doi:10.1016/j.jaci.2022.08.031

Singhal, D., Psaltis, A. J., Foreman, A., & Wormald, P. J. (2010). The impact of biofilms on outcomes after endoscopic sinus surgery. Am J Rhinol Allergy, 24(3), 169–174. doi:10.2500/ajra.2010.24.3462

Stanbery, A. G., Shuchi, S., Jakob von, M., Tait Wojno, E. D., & Ziegler, S. F. (2022). TSLP, IL-33, and IL-25: Not just for allergy and helminth infection. J Allergy Clin Immunol, 150(6), 1302–1313. doi:10.1016/j.jaci.2022.07.003

Takeda, K., Sakakibara, S., Yamashita, K., Motooka, D., Nakamura, S., El Hussien, M. A.,… Kikutani, H. (2019). Allergic conversion of protective mucosal immunity against nasal bacteria in patients with chronic rhinosinusitis with nasal polyposis. J Allergy Clin Immunol, 143(3), 1163–1175 e1115. doi:10.1016/j.jaci.2018.07.006

Teufelberger, A. R., Broker, B. M., Krysko, D. V., Bachert, C., & Krysko, O. (2019). Staphylococcus aureus Orchestrates Type 2 Airway Diseases. Trends Mol Med, 25(8), 696–707. doi:10.1016/j.molmed.2019.05.003

Tomassen, P., Vandeplas, G., Van Zele, T., Cardell, L. O., Arebro, J., Olze, H., … Bachert, C. (2016). Inflammatory endotypes of chronic rhinosinusitis based on cluster analysis of biomarkers. J Allergy Clin Immunol, 137(5), 1449–1456 e1444. doi:10.1016/j.jaci.2015.12.1324

Van Zele, T., Claeys, S., Gevaert, P., Van Maele, G., Holtappels, G., Van Cauwenberge, P., & Bachert, C. (2006). Differentiation of chronic sinus diseases by measurement of inflammatory mediators. Allergy, 61(11), 1280–1289. doi:10.1111/j.1398-9995.2006.01225.x

Van Zele, T., Gevaert, P., Watelet, J. B., Claeys, G., Holtappels, G., Claeys, C.,… Bachert, C. (2004). Staphylococcus aureus colonisation and IgE antibody formation to enterotoxins is increased in nasal polyposis. J Allergy Clin Immunol, 114(4), 981–983. doi:10.1016/j.jaci.2004.07.013

Vickery, T. W., Ramakrishnan, V. R., & Suh, J. D. (2019). The Role of Staphylococcus aureus in Patients with Chronic Sinusitis and Nasal Polyposis. Curr Allergy Asthma Rep, 19(4), 21. doi:10.1007/s11882-019-0853-7

Wang, X., Zhang, N., Bo, M., Holtappels, G., Zheng, M., Lou, H.,… Bachert, C. (2016). Diversity of T(H) cytokine profiles in patients with chronic rhinosinusitis: A multicenter study in Europe, Asia, and Oceania. J Allergy Clin Immunol, 138(5), 1344–1353. doi:10.1016/j.jaci.2016.05.041

Weigert, M., Schmidt, U., Haase, R., Sugawara, K., & Myers, G. (2020). Star-convex polyhedra for 3D object detection and segmentation in microscopy. Paper presented at the Proceedings of the IEEE/CVF Winter Conference on Applications of Computer Vision.

Wickham, H., Averick, M., Bryan, J., Chang, W., McGowan, L. D. A., François, R.,… Hester, J. (2019). Welcome to the Tidyverse. Journal of open source software, 4(43), 1686.

Wu, H., Moser, C., Wang, H. Z., Hoiby, N., & Song, Z. J. (2015). Strategies for combating bacterial biofilm infections. Int J Oral Sci, 7(1), 1–7. doi:10.1038/ijos.2014.65

Wu, T., Hu, E., Xu, S., Chen, M., Guo, P., Dai, Z.,… Yu, G. (2021). clusterProfiler 4.0: A universal enrichment tool for interpreting omics data. Innovation (Camb), 2(3), 100141. doi:10.1016/j.xinn.2021.100141

Yu, G., Li, F., Qin, Y., Bo, X., Wu, Y., & Wang, S. (2010). GOSemSim: an R package for measuring semantic similarity among GO terms and gene products. Bioinformatics, 26(7), 976–978. doi:10.1093/bioinformatics/btq064

